# Hypothalamic melanocortin-4 receptors on astrocytes mediate inflammation and body weight homeostasis

**DOI:** 10.1101/2022.12.01.518727

**Authors:** Nicole L. Eliason, Kevin Pham, Rohan Varshney, Jacob W. Farriester, Charles I. Lacy, Heather C. Rice, Michael C. Rudolph, Willard Freeman, Amanda L. Sharpe

## Abstract

World-wide, nearly 40% of the adult population is classified as clinically obese. Central inflammation is highly correlated with obesity and increases morbidity and deterioration of health. The hypothalamus is a brain region that governs many facets of energy homeostasis, and melanocortins in the hypothalamus both decrease feeding and increase metabolism via melanocortin-4 receptors (MC4Rs). Although MC4Rs are present on neurons and astrocytes (aMC4R) previous work has focused almost exclusively on the neuronal population with the contribution of aMC4R on these processes largely unknown. Our objective was to determine the effects of hypothalamic aMC4R deletion on central and peripheral inflammation, as well as feeding and body weight homeostasis. Adult MC4R fl/fl mice were microinjected with an astrocyte-specific promoter driving Cre-expression (AAV-GFAP-GFP-Cre) or AAV-control (AAV-GFAP-GFP; n=4-7/group/sex) to produce a hypothalamic knock-down of aMC4R (KD). Body weight and composition were monitored throughout the study, and indirect calorimetry was conducted at 1 and 4 weeks after AAV injection. Acquisition of operant self-administration of palatable food was also examined. Mice were euthanized 7-8 weeks post AAV injection and brain and tissue samples were collected. We observed a significant increase in body weight, feeding, and energy balance in the KD group compared to control group. Inflammation was significantly increased centrally in KD mice within the hypothalamus, but not peripherally within serum. Additionally, aMC4R KD mice trended towards an increased reward learning for palatable food. This is the first demonstration that hypothalamic aMC4R, independent of neuronal MC4R, is important in modulating inflammation as well as contributing to energy balance. These results provide an integral understanding of the aMC4R system that will provide the foundation for future studies investigating the role of aMC4R in various disease states.

## INTRODUCTION

Proper regulation of appetitive behaviors and metabolic processes are essential for maintenance of health within an organism. Modern social and environmental factors have, unfortunately, made the development of central inflammation strikingly more common. Chronic overnutrition, reduced physical activity and increased daily stressors all contribute to the acquisition and proliferation of inflammation-associated diseases.

The hypothalamus (HT) is one brain region that is uniquely connected to peripheral cues in both form and function. In form, the HT is a circumventricular organ, or a region within the brain where the blood-brain barrier becomes permeable (1). In function, the HT serves as the primary regulator of body homeostasis (2) within the brain with governing power over appetitive, thermal, metabolic and hormonal activities (3-5). This makes the HT a highly consequential target for neuroinflammation, and recent research has shown central inflammation can be observed prior to weight gain and drive acquisition of peripheral obesity (6). Furthermore, attenuation of inflammation exclusively in the HT is sufficient to significantly increase lifespan and reverse diet-induced insulin resistance (7,8).

One specific circuitry within the HT that has potent effects on body weight homeostasis and anti-inflammatory processes is the melanocortin (MC) system (9-11). While the contributions to anorexigenic behaviors are long-established, more recent studies have elucidated a role for MC’s in inflammation. Melanocortin-4 receptors (MC4R’s) have been specifically attributed for the central effects observed in both in-vitro and in-vivo models (12,13), and are expressed on both neuronal and glial cell types. Though MC4R is not specific to neurons, previous work has been completed predominantly in the neuronal population or in a non-specific manner. This gap within the literature originally arose from philosophical significance placed on neuronal functions, and has been perpetuated by prior inability to technically differentiate between cell-types.

While neurons drive a large portion of hypothalamic functions, recent emphasis within the MC system has shifted towards the astrocytic population. Astrocytes are glial cells that are uniquely positioned between the vasculature and neurons (14), making them highly-implicated in communicating circulating nutrient information to the central nervous system (15). Astrocytes essential for neuronal survival (16,17), and recent studies have identified critical roles for astrocytes in regulating inflammation, glucose uptake, and appetitive pathways within the HT (18,19). However, the contributions of astrocytes within the HT that express MC4R (aMC4R) have not been investigated. Our studies aimed to investigate the role of aMC4R exclusively within the hypothalamus on inflammation and body weight homeostasis through cell-type specific deletion.

## METHODS

### Animals

Adult (8-10 weeks) male (n=12) and female (n=11) MC4R fl/fl mice (Jackson Laboratory) were used for receptor characterization studies. Validation data was completed exclusively with male (n=15) MC4R fl/fl mice. All mice were maintained on ad libitum access to normal chow (PicoLab Rodent 20, LabDiet, MO) diet and were single-housed in a normal 12-h light/dark light cycle. Only mice tested in the operant study were housed in a reverse light-cycle, where they were given 4 days to acclimate prior to the start of the study. All procedures were approved by the University of Oklahoma Health Sciences Center Institutional Animal Care and Use Committee and were consistent with the best practices in *The Guide for Care and Use of Laboratory Animals*.

### Surgery

Mice were anesthetized with isoflurane (Isothesia NDC 11695-6776-2, Henry Schein, OH) at 2.0-2.3% for stereotaxic surgery. Briefly, mice were positioned into a stereotaxic frame (Kopf, Model 1900), and the surface of the skull was exposed through a single incision on the dorsal surface of the skull. Bregma was located, the skull was leveled, and holes were drilled at the proper coordinates (M/L = + 0.450/-0.500 mm; A/P = -1.000/-1.250 mm, relative to Bregma). Injection coordinates were slightly staggered both M/L and A/P to reduce skull fractures and ensure spread of the AAV injection throughout the HT. All injection coordinates were counterbalanced across groups to reduce experimental confounds. Injections were made using a syringe pump fitted with a 10-μl Hamilton syringe connected to injectors made from 33g hypodermic stainless-steel tubing (304SS Regular Wall; Component Supplies, TN) via PE20 tubing (BTPE-20, INSTECH, PA). Injectors were lowered into the drilled holes (D/V = -5.800 relative to skull surface at Bregma), and bilateral microinjections of either a Cre-expressing adeno-associated viral vector under an astrocyte-specific promoter (AAV5-GFAP-GFP-Cre; UNC Vector Core, NC) or control (AAV5-GFAP-GFP; UNC Vector Core, NC) viral vector were made. Injections were 400 nL in volume, and were administered at a rate of 0.2μl/min. Injections began at the lowest point (D/V = -5.800) and were raised to the level of PVN (D/V = -4.800) with 200 nL of volume remaining for the duration of the injection. Injectors were left in place for 5 minutes at the most dorsal position to facilitate diffusion away from the injection site before the injector was slowly retracted from the brain over 2-minutes. The skin incision was closed using cyanoacrylic adhesive. Before removal from anesthesia, mice were administered analgesia (meloxicam, 10 mg/kg; NDC: 11695-6936-2; Covetrus, ME) subcutaneously. The mice were checked regularly after surgery to monitor body weight and check the incision site. Mice were single housed and moved to a purified, normal fat diet (D11112201, 15% kcal from fat; Research Diets Inc, NJ) at least 24 hours after surgery and were allowed to recover for 1 week prior to their first indirect calorimetry session.

### Body Composition

Body composition analysis of mice used for indirect calorimetry studies was measured immediately 2 days post-injection, as well as immediately before and after indirect calorimetry measurements (1, 2, 4 and 5 weeks post injection). Fat and lean mass composition was measured using a non-invasive qMR: EchoMRI™ 4-in-1 Body Composition Analyzer (EchoMRI, TX).

### Indirect Calorimetry

Mice were transferred to Sable Systems International Promethion CAB-16 Environmental Control Cabinet (Sable Systems, NV) indirect calorimetry chambers at one and four to five weeks post-injection. Mice were single housed with ad libitum access to the purified diet (D11112201, 15% kcal from fat; Research Diets Inc, NJ) and water, as well as a housing module. Bodyweight, food-consumption, water intake, locomotion, oxygen consumption, and carbon dioxide production were measured during the testing period. Mice were given 3 days to habituate prior to 96 hours of continuous data collection for analysis.

### Operant Training

Approximately 7 weeks after surgery, male mice were introduced to the modular mouse operant chambers (Lafayette Instruments, Lafayette, IN). The operant sessions were controlled by and data were recorded on a computer with ABET software (Lafayette Instruments, Lafayette, IN). Each chamber was individually housed within a sound canceling exterior cabinet, accompanied by a fan to mask external noise. One wall of the chamber housed 2 nose-poke holes with a pellet receptacle centered between them. On the opposite wall mice had ad libitum access to a metal sipper nozzle connected to an inverted conical tube containing water. The feeder attached to the receptacle released a single 20-mg chocolate flavored fortified chow pellet (Cat. No. F05301; Bio Serv, NJ) upon completion of the assigned fixed ratio (FR) value. The side of the correct nose-poke for each mouse was counterbalanced across chambers to avoid bias from a side preference. Sessions began at 11 am, 1 hour after the dark portion of the light cycle began. Mice were trained to a fixed ratio (FR) value of 30, beginning with two consecutive days at an FR5 and doubling each subsequent session with acceptable activity. Mice were advanced to a higher FR ratio if they performed with an accuracy of 80% or higher the previous session. If the prior session activity was between 60 – 79% accurate they remained at the same FR value, and if accuracy below 60% the FR was reduced by an increment of 5. Acquisition of operant self-administration was defined as daily session accuracy greater than 90%. Activity throughout the acquisition period, as well as time to acquire, was collected for data analysis.

### Oral Glucose (OGTT) Tolerance Test

Male mice were fasted for 6 hours at the beginning of the dark cycle. blood glucose levels were measured using a Contour Next EZ blood glucose meter and Contour Next test strips. Blood glucose was measured 5 minutes before and 5, 10, 15, 20, 30, 45, 60, 90 and 120 minutes after oral gavage of 2 g dextrose/kg body weight with a 20% dextrose in water solution (Vanderbilt University Mouse Metabolic Phenotyping Center OGTT Protocol, dx.doi.org/10.17504/protocols.io.yz2fx8e).

### Perfusion and Tissue Preparation

At the conclusion of the study, tissues were collected and prepared for either cellular or histological analysis. For histology preparation, mice were deeply anesthetized with avertin (2, 2, 2-tribromoethanol; Sigma-Aldrich) prior to blood draw via cardiac puncture followed by transcardial perfusion with 10 mL of 10% sucrose in 1X Phosphate Buffered Saline (PBS), immediately followed by 10 ml 4% paraformaldehyde in 1X PBS. Brains were collected and post-fixed for 24 hours in 4% PFA, followed by 24-48 hours in 30% sucrose in 1X PBS. For cryosectioning, brains were embedded into OCT before sectioning (30-microns) on Cryostar NX50 Cryostat (Thermo Scientific, MA). Sections were placed into 24-well plates with 1X PBS for storage until immunostaining. For cell analysis preparation, mice were euthanized using cervical dislocation followed by decapitation. Brains and peripheral tissues (liver, fat and gastrocnemius) were immediately collected and placed into appropriate storage mediums for subsequent analysis.

### Blood Serum Analysis

Blood samples taken via cardiac puncture at euthanasia were allowed to clot for 2 hours at room temperature prior to centrifugation for 20 minutes at 4 degrees Celsius. Serum was carefully removed and aliquoted into fresh tubes for storage in -20 degrees Celsius until the day of analysis. Samples were processed per manufacturer’s protocol using a meso-scale discovery (MSD) multiplex immunoassay kit (V-PLEX Proinflammatory Panel 1 Mouse Kit; Cat# K15048D; Meso Scale Diagnostics, MD) consisting of TNF-alpha and IL-6 analytes only. Plates were read using an MSD Meso Quickplex SQ 120 instrument (Meso Scale Diagnostics, MD).

### Immunohistochemistry

Immediately after brains were sectioned, free-floating slices were manually selected for early to late hypothalamus. Within a 12-well plate, tissue was blocked using 2% Normal Donkey Serum (NDS; Cat No. 017-000-121; Jackson ImmunoResearch, PA) in 1X PBS. After a 30-minute blocking period and one 15-minute wash, 1.5 mL/well of primary antibody mix for GFAP (1:1000; Cat No. 287 004; EnCor Biotechnology, FL) and MC4R (1:250; Cat No. AMR-024; Alomone Labs, PA) in 1X PBS + 0.3% triton-X and 2% NDS was applied. Primary antibodies incubated for 6 hours at room-temperature before being moved to 4° C for approximately 48 hours. After incubation, slices were washed in 1X PBS for 15 minutes for a total of four washes. Secondary antibodies for rabbit (AF647; Ref No. A32795, Invitrogen, MA) and chicken (RRX; Cat No. 703-605-155; Jackson ImmunoResearch, PA) were diluted in 1X PBS and then added to the well-plates and incubated at room temperature for 2-hours before washing the sections 4 times in 1X PBS. After washes were complete, slices were mounted onto gelatin-coated Superfrost slides (Cat. No. 12-550-143; Fisher Scientific) and coverslipped using Invitrogen Prolong Diamond antifade mountant (P36961; Invitrogen, MA) and glass coverslips (Cat No. 12545F; Fisher Scientific, NH). Mountant was allowed to set on slides overnight to reduce photo-bleaching. Staining and mounting protocols were completed in an identical manner with the substitution of Iba1 (1:5000; RPCA-IBA1; EnCor Biotechnology, FL) as the sole primary antibody and rabbit (AF647; Ref No. A32795; Invitrogen, MA) as the secondary antibody for neuroinflammation analysis.

### Imaging

Slides for validation and inflammation studies were imaged using a Leica M205-MFC 2D/3D THUNDER large specimen imaging system microscope (Leica Microsystems, Buffalo Grove, IL) with the appropriate filters for the applied fluorophores. All images being compared across groups were taken at identical magnification and exposure. Images were taken using LAS X software (Leica Microsystems, Buffalo Grove, IL) and saved to a personal drive for subsequent analysis. High magnification images were taken with a Nikon Super-Resolution Spinning Disk Confocal microscope (Nikon Inc, NY).

### Statistical Analysis

Analysis of data were conducted with GraphPad Prism 9 (GraphPad Software, LLC). t-tests and ANOVAs were used appropriately as indicated in the results section. Significance was defined as alpha < 0.05.

## RESULTS

### Validation of adeno-associated virus specificity under a GFAP promoter in an MC4R fl/fl mouse model

Previous attempts at utilizing a GFAP promoter on an adeno-associated virus (AAV) to drive the expression of Cre have raised concerns about cell-type specificity under this particular promoter. To address this issue, our lab completed validation of the AAV-GFAP-GFP-Cre vector that we used to generate our astrocyte knockdown (aKD) model. We tested both serotype 5 (V5 aKD, UNC Vector Core) and PHP.eB (PHP.eB aKD, Addgene #10550) versions of this vector against control and neuronal knockdown (nKD, AAV-hSyn-Cre PHP.eB, Addgene #105540) viruses. Control and nKD designated minimal and maximal effects of MC4R deletion as knocking down neuronal MC4R should generate the most significant effects on phenotype. Body weight and immunostaining were used as physiological measures of viral specificity and MACSQuant flow cytometry analysis was used to quantify viral transfection and specificity. Weight gain is characteristic of MC4R deletion and therefore was measured over the 12-day period after viral injection. We observed that the PHP.eB aKD group, though still significantly lower, more closely resembled the final body weight (Figure 1A) and percent change in body weight from pre-injection weight (Figure 1B) of the nKD group than controls. Furthermore, the V5 aKD mice did not significantly differ in final body weight (Figure 1A) and gained weight in a similar trajectory to the controls (Figure 1C). Though the V5 aKD group did have a significantly higher percent change in body weight compared to controls (Figure 1B), these results suggest a higher degree of viral specificity in the V5 aKD vector given its similarities to the control group rather than the nKD.

**Figure 1.**
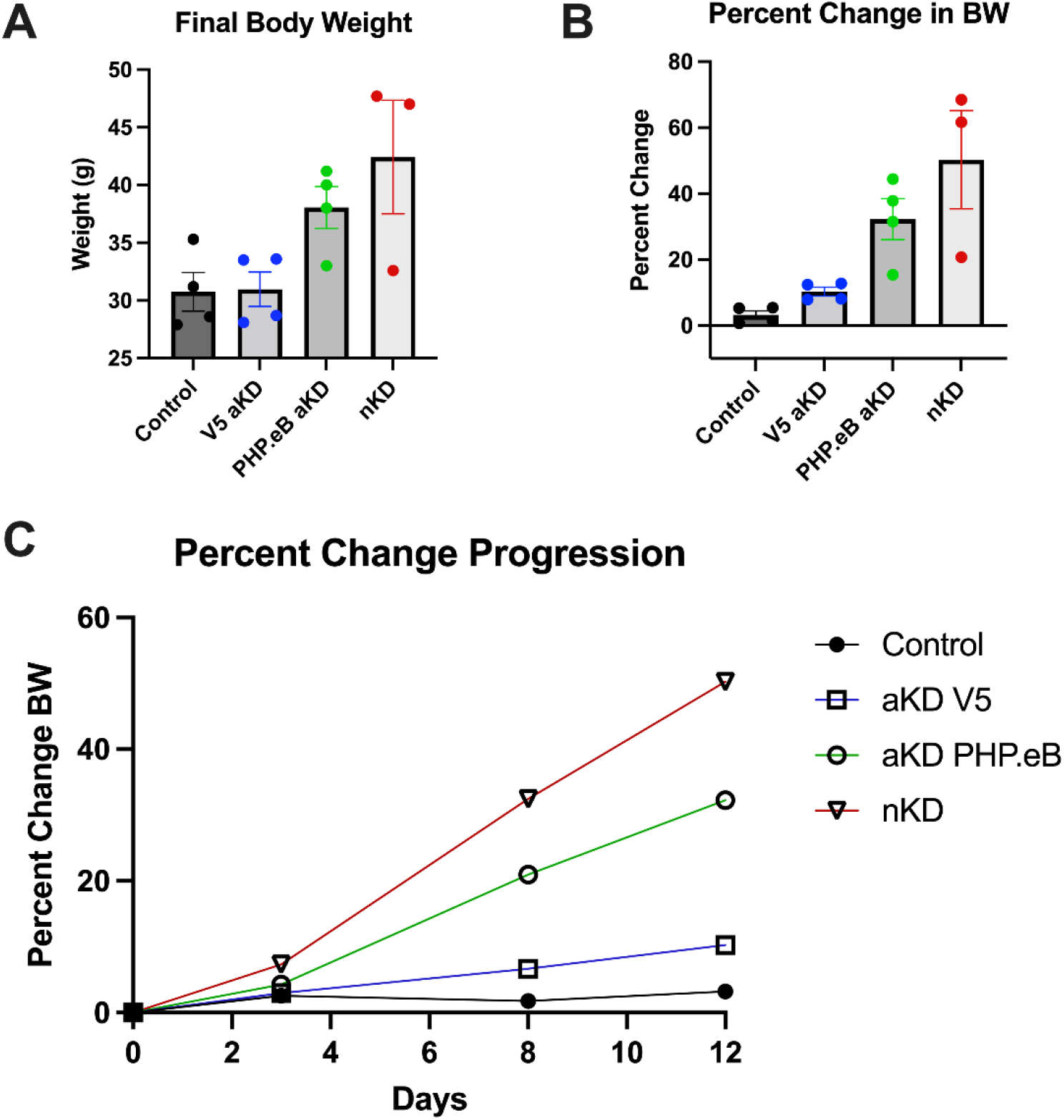
Direct comparison of effects of adeno-associated viral vectors on weight gain in MC4R fl/fl male mice. Mice were administered identical injections of control, GFAP-Cre serotype 5 (V5 aKD), GFAP-Cre PHP.eB (PHP.eB aKD) and hSyn-Cre PHP.eB (nKD) adeno-associated viral vectors. Body weight analysis over a 12-day period reveals that V5 aKD highly resembles total body weight (Panel A) of the control group *P=0.9324. This trend was conserved when analyzing percent change in total body weight (Panel B) as well as the trajectory of weight gain throughout the test window (Panel C) when compared to the nKD group. Furthermore, mice injected with the PHP.eB aKD displayed weight gain similar to the nKD group *P=0.3882, and significantly differed from controls in both total body weight *P=0.0252 and percent change in body weight *P=0.0037.

To determine the extent of receptor knockdown and localization between GFP (AAV transduction reporter), GFAP and MC4R, cryotome sectioned slices were stained for MC4R and GFAP. We observed severe knockdown of MC4R in the PHP.eB aKD group within the paraventricular nucleus of the hypothalamus (PVN) that mirrored MC4R signal reduction in the nKD group (Figure 2A). These results indicate that the PHP.eB aKD virus non-specifically infects cells which replicate the MC4R deletion pattern observed in the nKD group. Additionally, MC4R expression in the V5 aKD group was modestly reduced when compared to controls (Figure 2A) while retaining a healthy population within the PVN and indicating a greater degree of viral specificity than the PHP.eB aKD.

**Figure 2.**
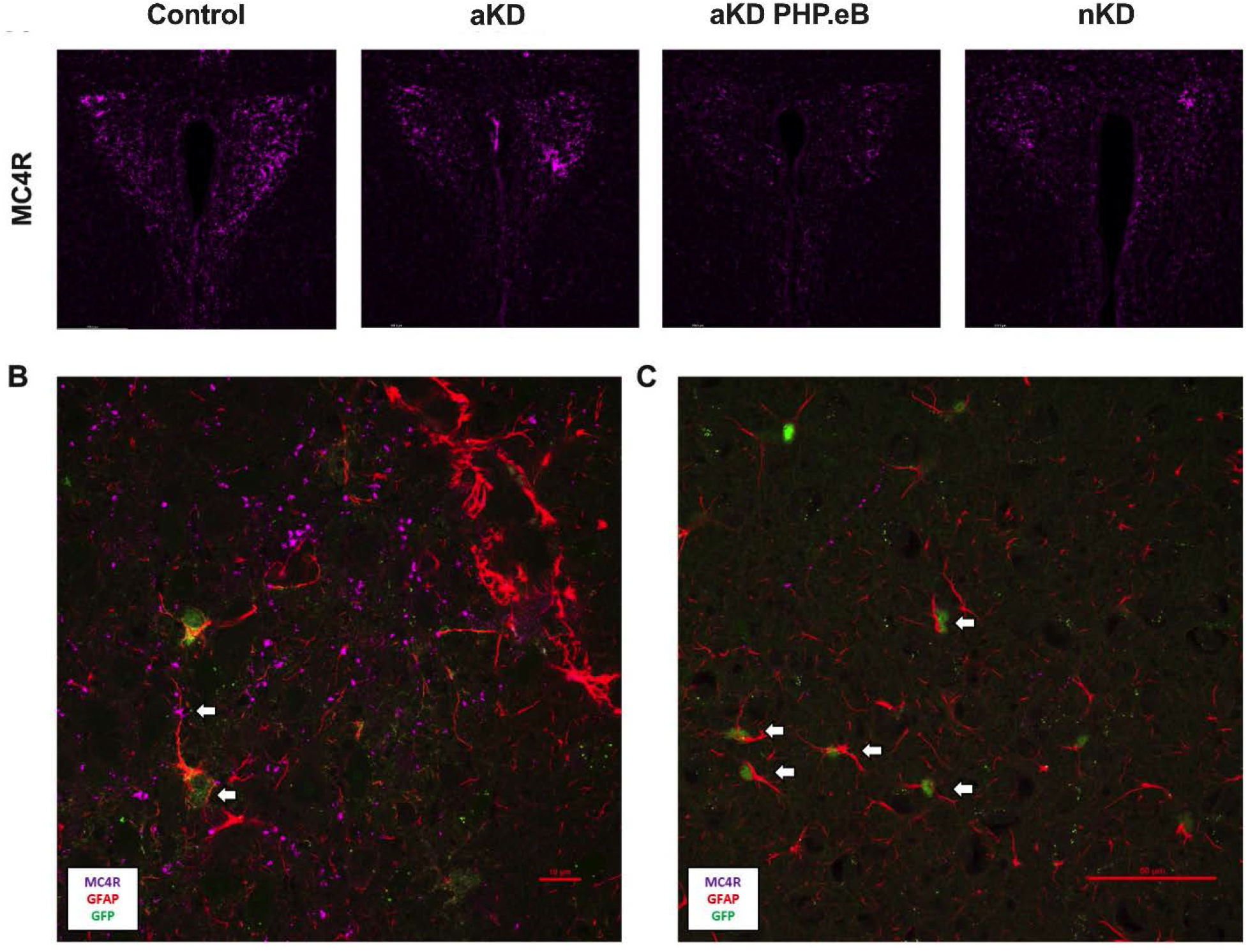
Comparison and localization of MC4R immunostaining in the PVN of AAV-injected mice. Immunostaining for MC4R in the PVN across treatment groups (Panel A) shows the progression of aMC4R KD from control, aMC4R specific, aMC4R nonspecific and specific nMC4R viral vectors. Confocal imaging of control (AAV-GFAP-GFP) mice demonstrated colocalization of GFP, GFAP and MC4R in the cell body and along the dendrite (Panel B), as well as non-colocalization of MC4R with GFP positive astrocytes in experimental (AAV-GFAP-GFP-Cre) injected mice (Panel C).

Lastly, a small cohort of mice (n=2) were injected with the V5 aKD virus and brain hypothalami were collected for MACSQuant analysis. Evaluation of the pooled brain sample revealed a very small transfection rate of cells within the hypothalamus (Figure 3B) and a high degree of specificity when colocalizing with ACSA2 (antibody for astrocytes; Figure 3D). These results suggest that effects mediated by this viral knockdown are not only highly specific but are mediated by a small fraction of the overall astrocyte population in the hypothalamus.

**Figure 3.**
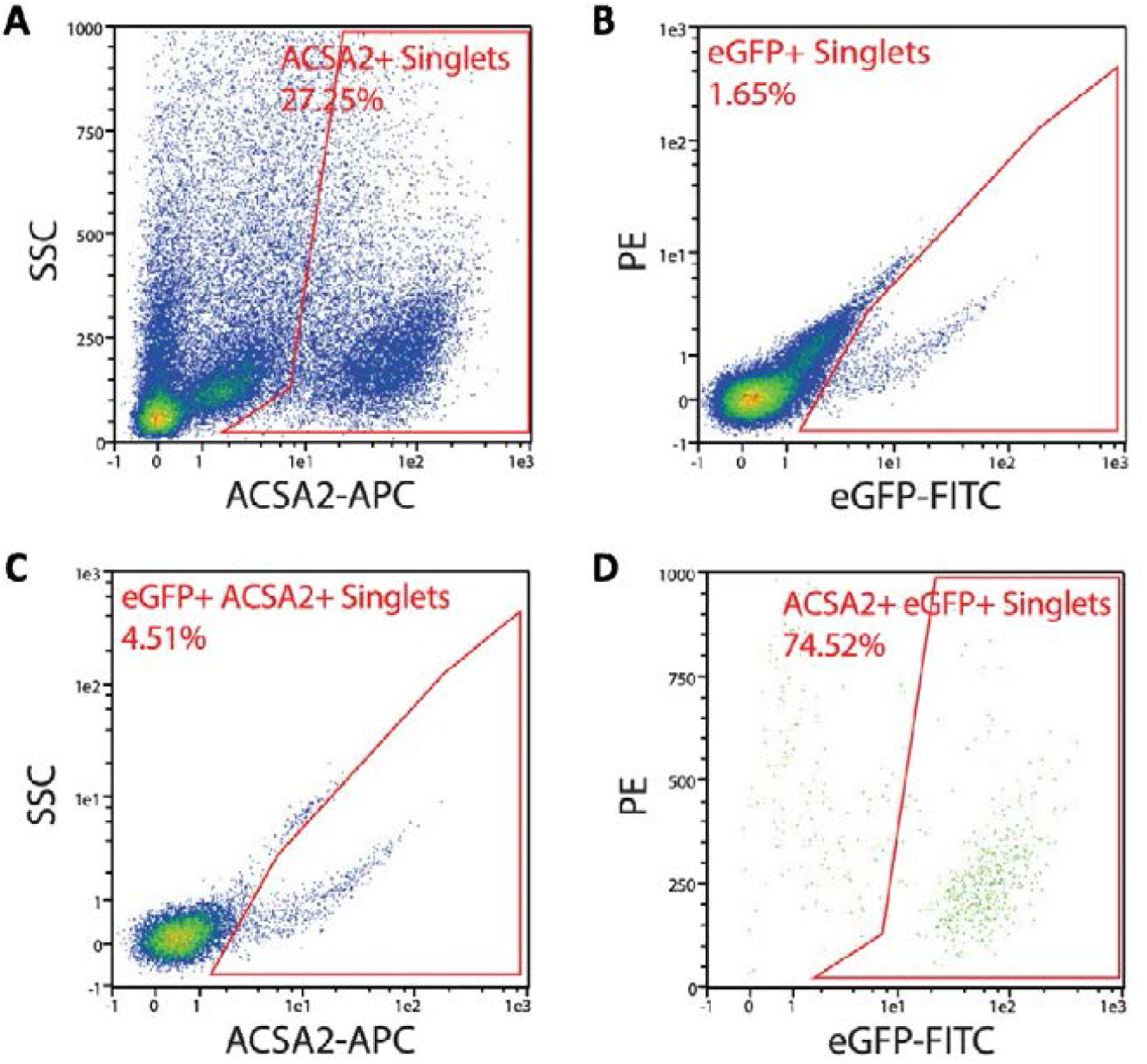
MACS-Quant analysis of isolated hypothalamic cells from V5 aKD injected mice. Analysis of unsorted cells stained with ACSA2 (astrocytic marker) shows that 27.25% of cells isolated from the hypothalamus are ACSA2 positive (Panel A), while only 1.65% of the total cell count are eGFP positive (Panel B) from successful viral transfection. Furthermore, 4.51% of the total ACSA2 positive population was also eGFP positive (Panel C). Lastly, the V5 aKD displayed 74.52% specificity when colocalizing with the ACSA2 positive cell population (Panel D).

### Hypothalamic deletion of aMC4R significantly increased the presence of neuroinflammatory markers in the hypothalamus

Mice used for the viral validation studies were also used to determine the presence of central inflammation via immunostaining. GFAP presence and proliferation, as well as increases in Iba1 (microglial activation) staining are accepted indicators of neuroinflammation (20,21). We observed a visual increase (Figure 4A) as well as a statistically significant increase in both GFAP (Figure 4B) and Iba1 (Figure 4C) cell density within the PVN of aKD mice when compared to controls. These results indicate that deletion of MC4R on astrocytes is sufficient to induce central inflammation within the hypothalamus.

**Figure 4.**
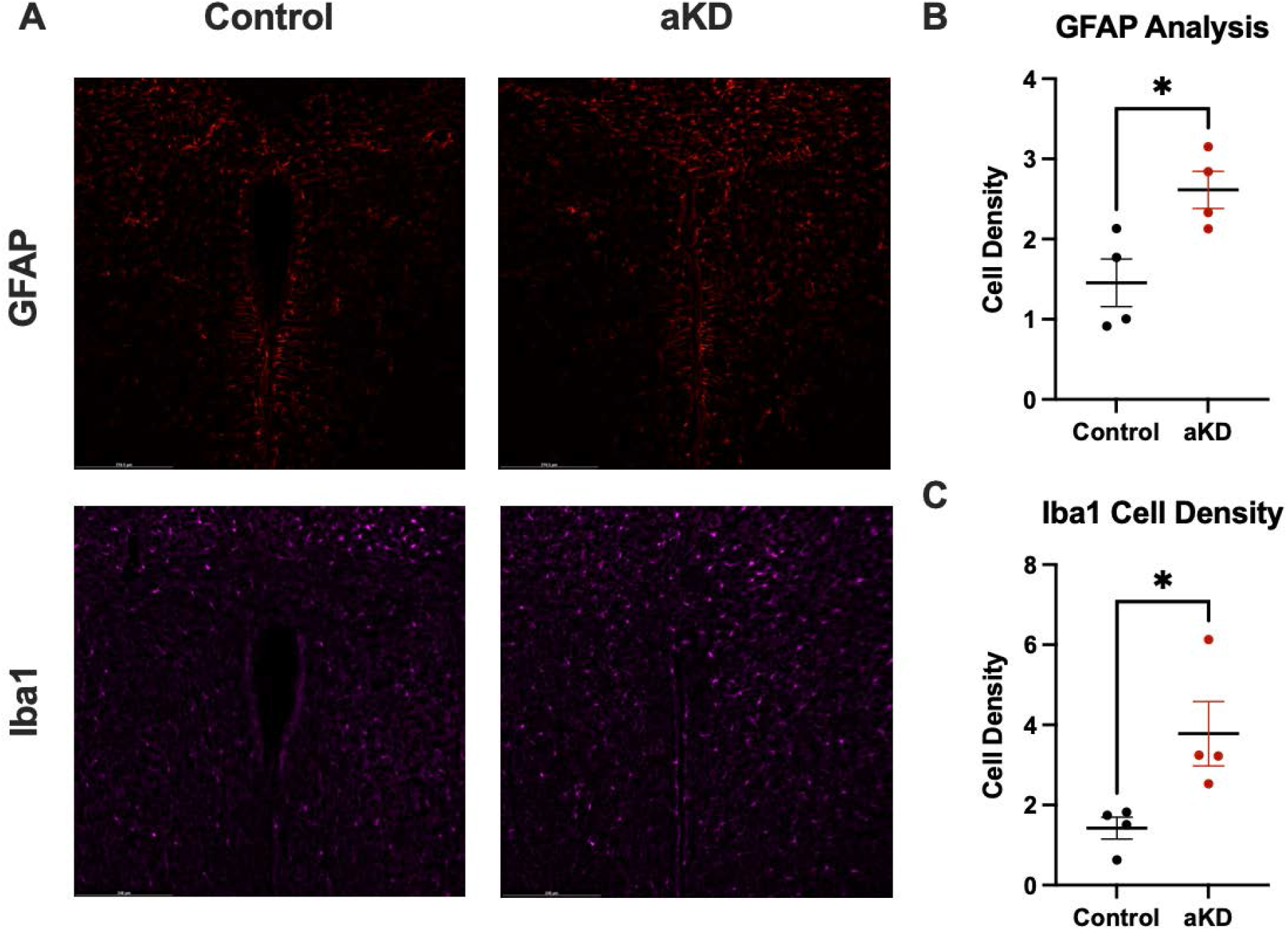
Effects of aMC4R deletion in the hypothalamus on neuroinflammation within the PVN using immunostaining. Brain slices from control and V5 aKD injected mice were stained for GFAP and Iba1 within the PVN. Quantification of these images for cell density completed using ImageJ ITCN analysis revealed significantly higher density of GFAP *P=0.0218 and Iba1 *P=0.0318 in aMC4R KD mice when compared to controls (Panel B,C).

### Markers of peripheral inflammation in blood serum are not significantly elevated in an aMC4R KD mouse model

Increased circulation of inflammatory markers is one of many biological components that acts to perpetuate the inflammation-obesity axis (22). Studies have found significantly increased serum levels of proinflammatory cytokines TNF-alpha and IL-6 in obese subjects (23). To determine whether the central inflammation induced by aMC4R deletion in the hypothalamus is driving similar inflammation in the periphery, we analyzed blood serum samples from mice at two time-points after AAV injection. TNF-alpha and IL-6 have been established to increase in blood serum within inflamed and obese mice (24) and are accepted indicators of inflammation within the periphery. We collected blood samples via direct cardiac puncture from male mice 3 weeks post-injection and male and female mice 8 weeks post-injection. We did not observe a significant difference in TNF-alpha concentration between control and aKD mice at either time point (Figure 5A, 5C). Furthermore, IL-6 was slightly elevated in the aKD group at both timepoints, but this increase was not found to be significant. While there was no effect of treatment on either serum inflammatory markers, there was a highly significant effect of time on concentration of both markers across groups. Given that the effect of time is not significantly different between treatment groups, we can conclude that the elevation in serum concentration with time is not attributed to the deletion of aMC4R. These results suggest that while aMC4R signaling disruption is sufficient to induce neuroinflammation, inflammation within the periphery is not significantly affected by hypothalamic aMC4R deletion.

**Figure 5.**
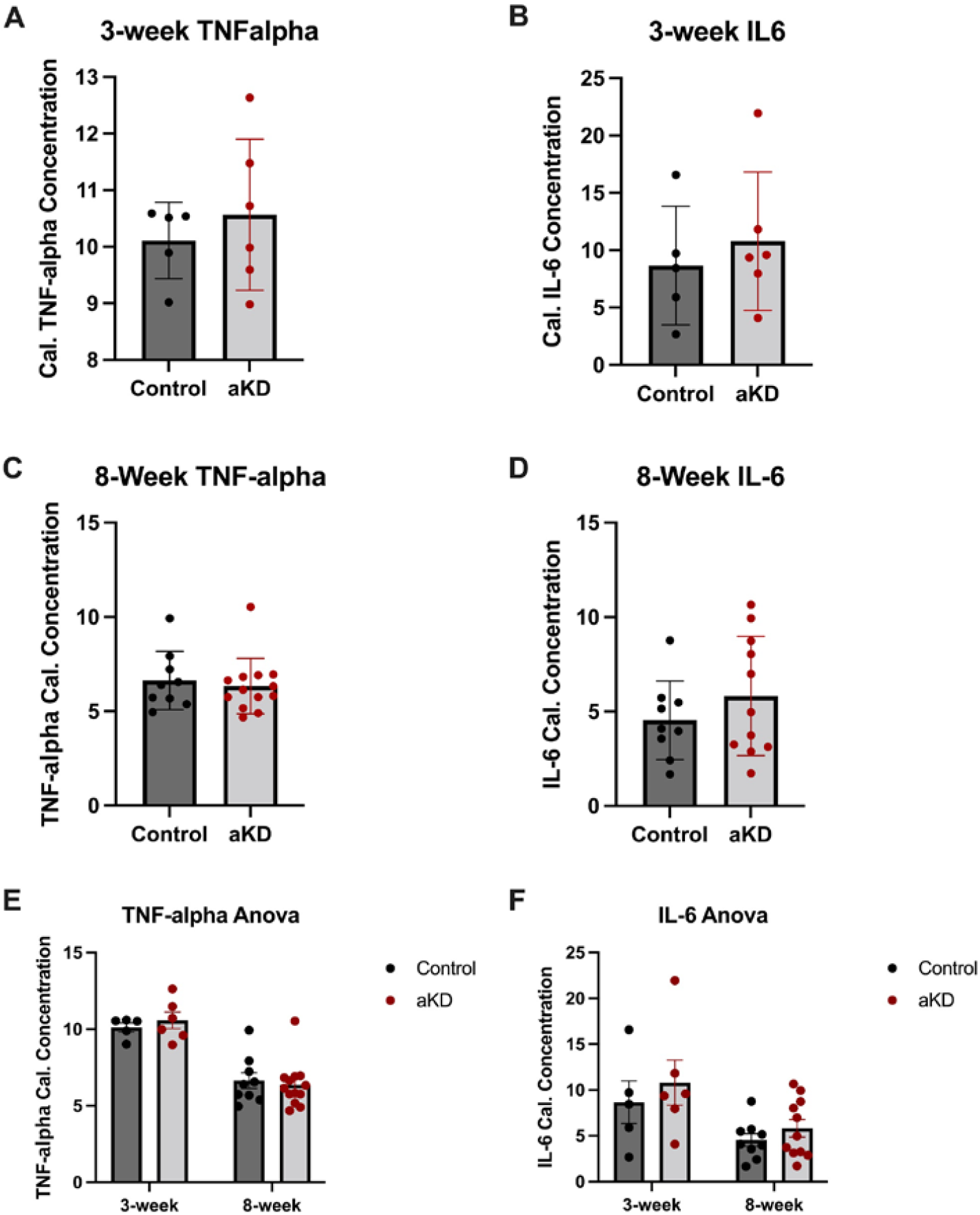
Serum analysis for markers of peripheral inflammation. Serum analysis of control and experimental injected mice showed no significant difference between control and experimental treated mice in TNF-alpha or IL-6 concentration at either time-point (Panel A-D). Analysis revealed a significant effect of time on both TNF-alpha *P=<0.0001 and IL-6 *P=0.0051 concentration across groups.

### Hypothalamic aMC4R KD mice had increased body weight and fat mass with no change in lean mass

While neuronal MC4R’s are well established to modulate body weight homeostasis (11), the contributions of the astrocytic population had not been investigated. The intermediary nature of the astrocyte population with neurocircuitry within the hypothalamus (25) would suggest that MC4R contributions would be largely anti-inflammatory. Surprisingly, we observed a significant increase in percent change of body weight within the aMC4R KD group when compared to controls (Figure 6A) that was conserved across both sexes. Echo MRI qMR body composition analysis was taken at multiple time-points to further investigate changes in lean and fat mass throughout the 5-week observation period post-injection (PI). In consensus with our body weight data, we observed a significant increase in percent change fat mass (Figure 6C) in the aMC4R KD group with no difference in percent change lean mass (Figure 6B) when compared to controls. While the obesigenic phenotype was conserved across sexes, we observed a bolstered effect within the male mice which had a highly significant increase in total fat mass gained (Figure 6D) at 5-weeks PI compared to BL.

**Figure 6.**
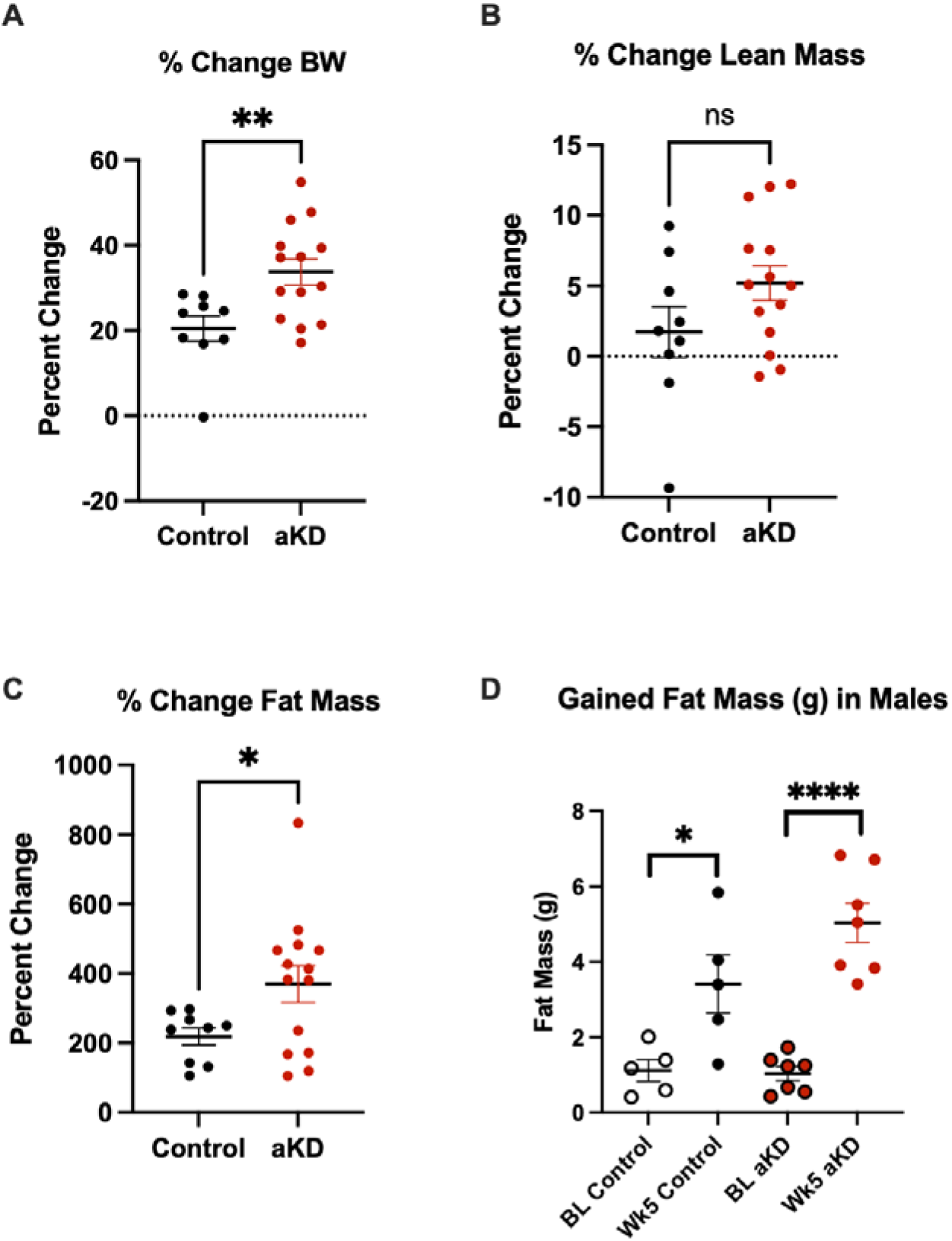
Comparison of body weight and composition among control and aMC4R KD mice. Percent change in body weight from body weight (g) on the day of surgical injections to the end of observation (5-weeks PI; Panel A) was significantly increased in experimentally injected aMC4R KD mice*P=0.0072 when compared to controls. Baseline (BL) body composition analysis was completed at 1-week PI and final values were collected at 5-weeks PI. Percent change in lean mass (g) was not different between the groups (Panel B), while percent change from BL of fat mass (g) was significantly increased *P=0.0403 in the experimental group (Panel C). We observed a significant increase in fat mass (g) across treatment groups from BL to 5-weeks PI (Panel D) in male mice with a more highly-significant increase in the experimental group *P=<0.0001 than controls *P=0.0230.

### aMC4R KD within the hypothalamus increases energy balance through hyperphagia and pedestrian locomotion

MC4R signaling pathways play a multi-faceted role in regulating energy balance. Stimulating MC4R’s within the brain has been shown to increase energy expenditure, locomotor activity and anorexigenic behaviors (26). We used promethion indirect calorimetry to investigate the effects of aMC4R deletion in the hypothalamus on these measures. Male and female mice were housed at two timepoints (1 and 4 weeks) post-injection and monitored continuously for 96 hours after acclimation. Dissimilar to previous studies investigating MC4R activity, we did not observe a significant difference in energy expenditure (Figure 7A) between the groups. However, we did observe a significant increase in energy balance within the aMC4R KD group (Figure 7D) that appears to be attributed to changes in food-intake and locomotor activity. Specifically, aMC4R KD mice were significantly hyperphagic when compared to controls (Figure 7C), and food consumption within the experimental group was consistent throughout both test periods. In addition, aMC4R mice had a significant reduction in locomotor activity (Figure 7B) that continued to decline over the observation period. These findings suggest an integral role for hypothalamic aMC4R in maintaining a healthy energy balance.

**Figure 7.**
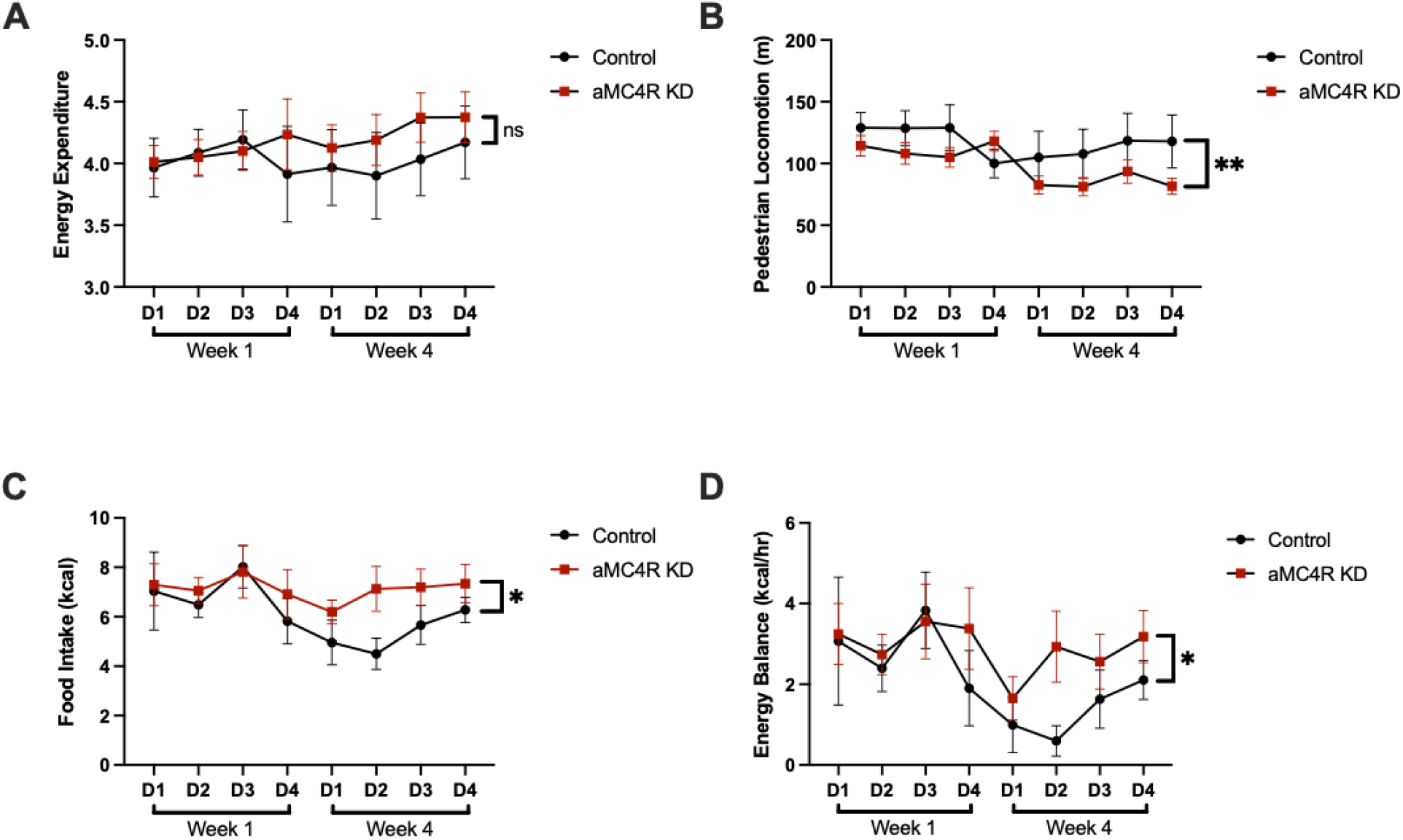
Investigation of energy balance shift using indirect calorimetry. Daily total measures for male and female mice during the dark window were collected to investigate contributions to energy balance within control and aMC4R KD mice. Energy expenditure (kcal/hr) was not significantly different between treatment groups (Panel A) and was consistent throughout both sessions. Pedestrian locomotion (m) was significantly reduced within the aMC4R KD *P=0.0049 (Panel B) group when compared to controls. aMC4R KD mice displayed a significantly hyperphagic phenotype *P=0.0219 which increased over time compared to the control treated mice (Panel C). The combination of reduced locomotor activity and increased food intake resulted in a significantly more positive energy balance *P=0.0453 within the experimental treated mice (Panel D).

### Hypothalamic aMC4R KD reduces operant food consumption and behavioral acquisition

Previous work by our lab and others has established a critical role for the melanocortin system in acquisition of food-based operant behavior. Particularly, that stimulation of proopiomelanocortin (POMC) projections in the nucleus accumbens bolsters acquisition and accuracy compared to saline controls. Hypothalamic inflammation induced by MC4R deletion is suspected to dysregulate POMC signaling which could negatively impact behavioral acquisition. To investigate this question, we tested a cohort of male mice where we observed a significant decrease in activity (correct nosepokes) on day 1 of training (Figure 8A) for aMC4R KD mice that was modestly conserved throughout the remaining training period (Figure 8B). Interestingly, we did not find a significant reduction in accuracy of aMC4R KD mice on training day 1 (Figure 8C) paired with the reduction in activity. Lastly, we assessed the days to acquire for the cohort where acquisition is defined as completing and maintaining an accuracy of 90% or greater. While we did not find a significant difference between treatment groups (Figure 8D), the only mice that took longer than 4 days to acquire the task were within the experimental group. These data support a model in which hypothalamic inflammation, via aMC4R deletion, negatively impacts the capacity for POMC neurons to establish reward learning.

**Figure 8.**
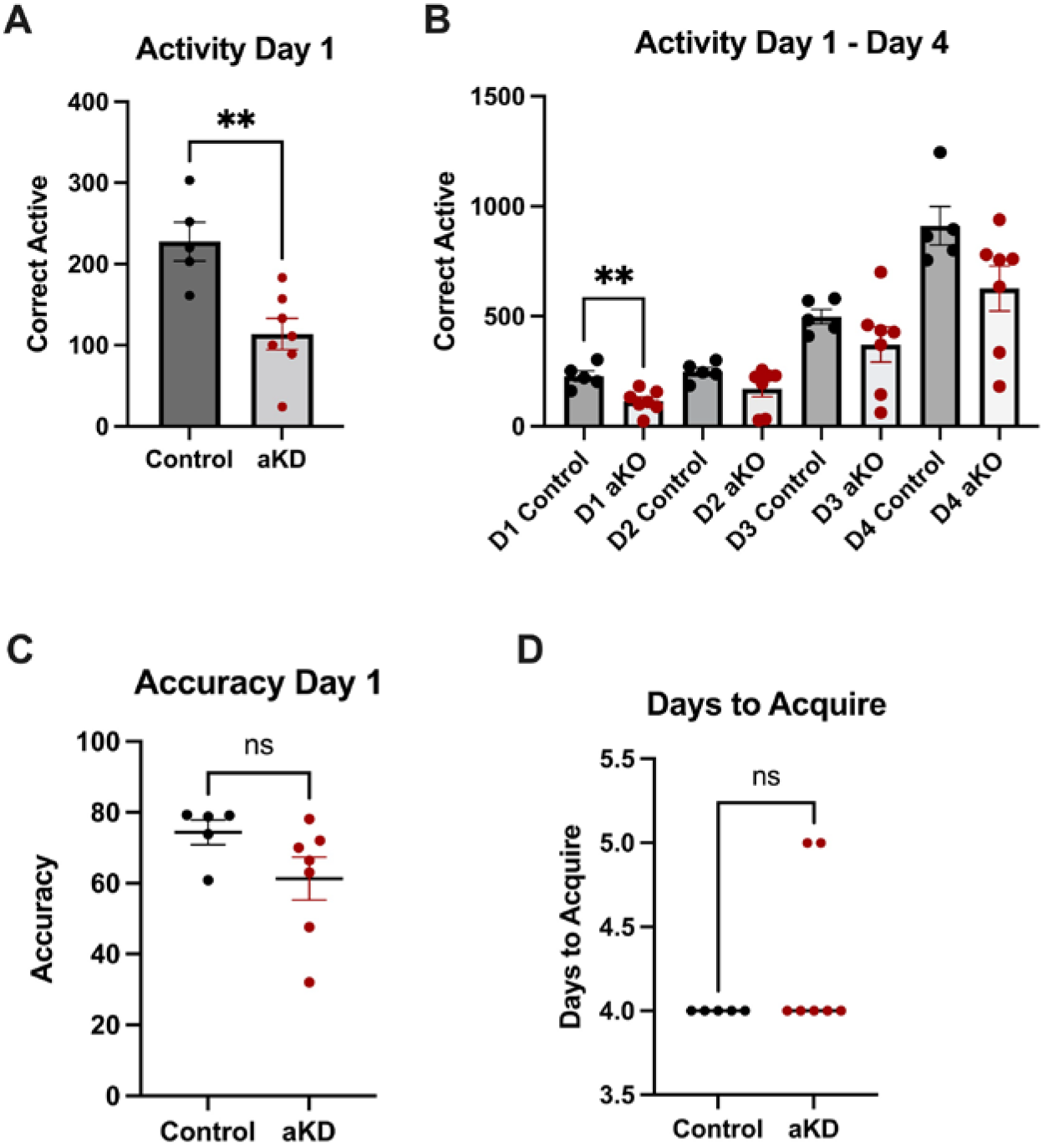
Food-based operant task acquisition and accuracy. Correct active nosepokes throughout the 3-hour session on Day 1 of operant training (Panel A) displayed significantly reduced activity in male aKD mice *P=0.0039 compared to controls. Activity within the aKD mice remained reduced throughout the 4-day training period (Panel B). Accuracy as percentage of correct nose pokes on Day 1 of training (Panel C) was not different between groups. Days to acquire were not significantly different between groups (Panel D). aKD mice were the only to display delayed behavior acquisition where acquisition was defined as accomplishing and maintaining an accuracy of 90% or greater.

## DISCUSSION

In this study we identified that hypothalamic aMC4R’s play an integral role in maintaining body weight homeostasis and modulating inflammation within the brain. This study was largely made possible by the novel combination of an MC4R fl/fl mouse line with a site-specific injection of an astrocyte specific viral vector. However, initial attempts at applying this technique using a PHP.eB version of this vector had proved problematic for cell-type specificity. Preliminary studies completed in our lab found that the PHP.eB virus generated an exaggerated phenotype in vivo and was histologically non-specific when using immunostaining. These initial results led us to rigorously validate the serotype 5 (AAV5) viral vector which was used throughout the study. In consensus with prior studies surrounding AAV5 specificity in CNS tissue (27), we determined that our virus had a high-degree of specificity when colocalizing with ACSA2. Given that the virus had a GFAP promoter, rather than ACSA2, our data are likely a conservative value. If MACSQuant was completed for colocalization with GFAP, the projected specificity should surpass the 75% marker reported in this study. Furthermore, immunostaining supported these findings, where we were unable to colocalize a GFP positive cell in a Cre-injected mouse with antibodies for MC4R. Future RNA-sequencing studies to investigate differential genetic and epigenetic modification will lend crucial insight to affected molecular pathways, though technical limitations surrounding cell-counts exist.

While previous work has focused on neuroinflammation upon the acquisition of obesity, recent studies have reiterated the preceding of inflammation to the development of obesity (6). These findings make maintenance of neuronal health and signaling paramount to prevention of obesity and chronic inflammation. Studies have implicated both IKKβ and NF-κB pathways as essential modulators of central inflammation, with particular regard to energy balance and obesity (8,10,28,29). Furthermore, MC4R activation on astrocytes has been shown to inhibit translocation of NF-κB to the nucleus (12), promoting an anti-inflammatory cellular environment. While our study did not investigate the cellular pathways mechanisms driving the observed phenotype, this evidence within the literature provides a rationale for further investigation into effects of aMC4R deletion on NF-κB and IKKβ pathways. Lastly, brain-derived neurotrophic factor (BDNF) modulates energy balance downstream of MC4R and suppressed expression of its receptor TrkB in the hypothalamus leads to hyperhagia-induced obesity (30). The relationship between BDNF and MC4R is hypothesized to occur between nMC4R and TrkB in the ventromedial hypothalamus, but warrants further investigation into potential direct/indirect effect of aMC4R activity within this system.

Serum sample analysis was completed to investigate concentrations of TNF-alpha and IL-6 within the periphery that are commonly associated with obesity (23). The presence of elevated inflammatory cytokines concentration is characteristically observed in diabetic patients (31), but has been linked to melanocortin modulated processes in traumatic brain inflammation (32). Given the obese phenotype we observed, the nonsignificant elevation in TNF-alpha and IL-6 was initially surprising. However, OGTT completed in the male mice revealed modestly improved glucose clearance in the experimental treated mice compared to controls, regardless of significantly elevated fasting blood glucose (Supplementary Figure 1A,1B). These results suggest that insulin sensitivity and efficacy is normal in these mice, and therefore we would not expect to see elevated TNF-alpha levels (33).

While a significant obese phenotype was not originally expected, the significant induction of inflammation within the hypothalamus upon deletion of aMC4R would constitute hyperphagia via disruption of neuronal signaling. Body composition was consistent with what has been observed in hyperphagia-induced obesity, though the weight gained as fat mass in the aMC4R KD group is more extreme than indirect calorimetry metrics would suggest. This is in part due to the lack of difference between groups when analyzing energy expenditure (EE). The observed reduction in pedestrian locomotion would suggest a decrease in EE of the aMC4R KD mice. However, the aMC4R mice were reported to have higher continuous rate of EE than the controls. These conflicting results may be in part to a few confounding variables. The first of which is CalR, the software used to obtain these data has limitations when accounting for individual animal body weight and its progression over time (34). While we are able to manually input initial body weight and composition data to generate these data, only one static value is used throughout the duration of the IDC run. Secondly, the central melanocortin system is documented to play a role in adaptive thermogenesis of brown adipose tissue (BAT, 35). Activation of BAT to generate heat will directly increase EE in a manner that is dependent on BAT mass (36). Given that BAT recruitment and mass is decreased in obesity (37), and the experimental design to disrupt central melanocortin signaling, the data surrounding EE may not be an accurate measure of the actual animal physiology.

Operant behavioral testing to investigate food-based reward learning supported prior work in our lab that hypothalamic POMC projections are essential for behavior acquisition in mice. The reduction in food consumption within the aMC4R KD group is likely attributed to a hindered ability to assign rewarding properties to a novel stimulus, rather than an inverse response to diminished anorexigenic melanocortin activity. The impact of these results are limited by the study being only completed in males, a shortened training period to an FR20 rather than an FR30, and a small sample size. These data provide a preliminary insight into what could be a more significant interaction if replicated in a larger, inclusive study.

Perhaps the largest question that remains upon the conclusion of this study is whether aMC4R modulates the affected processes directly through downstream signaling, or indirectly by maintaining an anti-inflammatory tone within the hypothalamus. There is significant evidence in previous literature and throughout our study to suggest that aMC4R KD effects are mediated through both direct and indirect interactions. The results from this study are the first to demonstrate an endogenous tonic role of melanocortins in preventing inflammation in the hypothalamus. This presents a possible link between loss of endogenous melanocortin activity at aMC4R causing the development of inflammation that then leads to hyperphagia and obesity. The results from this study are limited by the small population of cells that were affected (∼1.65% of total hypothalamic cells, Fig. 3B) and the fact that GFAP, the promoter used, is only present in a subset of astrocytes. Future studies should be conducted to delete aMC4R in a more pan-astrocytic manner to give a more thorough view of the role of aMC4R in inflammation and metabolic homeostasis.

## Acknowledgements

The authors would like to thank the University of Oklahoma Geroscience MACI core for access to inverted and confocal microscopy as well as the Oklahoma Medical Research Foundation Imaging Core for access to histology equipment. The authors would like to acknowledge funding for this research generously provided by the following grants: National Institutes of health grant P20GM12528 (sub-project 5338 to ALS and sub-project 5990 to HCR), Presbyterian Health Foundation Seed Grants (ALS) and OUHSC College of Pharmacy Seed Grant (ALS).

## References

1. Gross PM. Circumventricular organ capillaries. Prog Brain Res. 1992;91:219–33. PMID: 1410407.

2. Date Y, Ueta Y, Yamashita H, Yamaguchi H, Matsukura S, Kangawa K, Sakurai T, Yanagisawa M, Nakazato M. Orexins, orexigenic hypothalamic peptides, interact with autonomic, neuroendocrine and neuroregulatory systems. Proc Natl Acad Sci U S A. 1999 Jan 19;96(2):748–53. doi: 10.1073/pnas.96.2.748. PMID: 9892705; PMCID: PMC15208.

3. Anand BK, Brobeck JR. Localization of a “feeding center” in the hypothalamus of the rat. Proceedings of the society for experimental biology and Medicine. 1951 Jun;77(2):323–5. doi: 10.3181/00379727-77-18766. PMID: 14854036.

4. Daniel PM. Anatomy of the hypothalamus and pituitary gland. J Clin Pathol Suppl (Assoc Clin Pathol). 1976;7:1–7. doi: 10.1136/jcp.s1-7.1.1. PMID: 1073162; PMCID: PMC1436118.

5. Boulant JA. Hypothalamic mechanisms in thermoregulation. Fed Proc. 1981 Dec;40(14):2843–50. PMID: 6273235.

6. Thaler, J. P., Yi, C.-X., Schur, E. A., Guyenet, S. J., Hwang, B. H., Dietrich, M. O., Zhao, X., Sarruf, D. A., Izgur, V., Maravilla, K. R., Nguyen, H. T., Fischer, J. D., Matsen, M. E., Wisse, B. E., Morton, G. J., Horvath, T. L., Baskin, D. G., Tschöp, M. H., & Schwartz, M. W. (2012). Obesity is associated with hypothalamic injury in rodents and humans. The Journal of Clinical Investigation, 122(1), 153–162. https://doi.org/10.1172/JCI59660

7. Milanski M, Arruda AP, Coope A, Ignacio-Souza LM, Nunez CE, Roman EA, Romanatto T, Pascoal LB, Caricilli AM, Torsoni MA, Prada PO, Saad MJ, Velloso LA. Inhibition of hypothalamic inflammation reverses diet-induced insulin resistance in the liver. Diabetes. 2012 Jun;61(6):1455–62. doi: 10.2337/db11-0390. Epub 2012 Apr 20. PMID: 22522614; PMCID: PMC3357298.

8. Zhang G, Li J, Purkayastha S, Tang Y, Zhang H, Yin Y, Li B, Liu G, Cai D. Hypothalamic programming of systemic ageing involving IKK-β, NF-κB and GnRH. Nature. 2013 May 9;497(7448):211–6. doi: 10.1038/nature12143. Epub 2013 May 1. PMID: 23636330; PMCID: PMC3756938.

9. Benoit SC, Air EL, Coolen LM, Strauss R, Jackman A, Clegg DJ, Seeley RJ, Woods SC. The catabolic action of insulin in the brain is mediated by melanocortins. J Neurosci. 2002 Oct 15;22(20):9048–52. doi: 10.1523/JNEUROSCI.22-20-09048.2002. PMID: 12388611; PMCID: PMC6757684.

10. Ichiyama T, Sakai T, Catania A, Barsh GS, Furukawa S, Lipton JM. Inhibition of peripheral NF-kappaB activation by central action of alpha-melanocyte-stimulating hormone. J Neuroimmunol. 1999 Oct 29;99(2):211–7. doi: 10.1016/s0165-5728(99)00122-8. PMID: 10505977.

11. Cowley MA, Smart JL, Rubinstein M, Cerdán MG, Diano S, Horvath TL, Cone RD, Low MJ. Leptin activates anorexigenic POMC neurons through a neural network in the arcuate nucleus. Nature. 2001 May 24;411(6836):480–4. doi: 10.1038/35078085. PMID: 11373681.

12. Kamermans A, Verhoeven T, van Het Hof B, Koning JJ, Borghuis L, Witte M, van Horssen J, de Vries HE, Rijnsburger M. Setmelanotide, a Novel, Selective Melanocortin Receptor-4 Agonist Exerts Anti-inflammatory Actions in Astrocytes and Promotes an Anti-inflammatory Macrophage Phenotype. Front Immunol. 2019 Oct 4;10:2312. doi: 10.3389/fimmu.2019.02312. PMID: 31636637; PMCID: PMC6788433.

13. Muceniece R, Zvejniece L, Vilskersts R, Liepinsh E, Baumane L, Kalvinsh I, Wikberg JE, Dambrova M. Functional evaluation of THIQ, a melanocortin 4 receptor agonist, in models of food intake and inflammation. Basic Clin Pharmacol Toxicol. 2007 Dec;101(6):416–20. doi: 10.1111/j.1742-7843.2007.00133.x. PMID: 18028105.

14. Kacem K, Lacombe P, Seylaz J, Bonvento G. Structural organization of the perivascular astrocyte endfeet and their relationship with the endothelial glucose transporter: a confocal microscopy study. Glia. 1998 May;23(1):1–10. PMID: 9562180.

15. Gruber T, Pan C, Contreras RE, Wiedemann T, Morgan DA, Skowronski AA, Lefort S, De Bernardis Murat C, Le Thuc O, Legutko B, Ruiz-Ojeda FJ, Fuente-Fernández M, García-Villalón AL, González-Hedström D, Huber M, Szigeti-Buck K, Müller TD, Ussar S, Pfluger P, Woods SC, Ertürk A, LeDuc CA, Rahmouni K, Granado M, Horvath TL, Tschöp MH, García-Cáceres C. Obesity-associated hyperleptinemia alters the gliovascular interface of the hypothalamus to promote hypertension. Cell Metab. 2021 Jun 1;33(6):1155-1170.e10. doi: 10.1016/j.cmet.2021.04.007. Epub 2021 May 4. PMID: 33951475; PMCID: PMC8183500.

16. Mitoma J, Furuya S, Hirabayashi Y. A novel metabolic communication between neurons and astrocytes: non-essential amino acid L-serine released from astrocytes is essential for developing hippocampal neurons. Neurosci Res. 1998 Feb;30(2):195–9. doi: 10.1016/s0168-0102(97)00113-2. PMID: 9579653.

17. Allen NJ. Astrocyte regulation of synaptic behavior. Annu Rev Cell Dev Biol. 2014;30:439–63. doi: 10.1146/annurev-cellbio-100913-013053. PMID: 25288116.

18. Kim JG, Suyama S, Koch M, Jin S, Argente-Arizon P, Argente J, Liu ZW, Zimmer MR, Jeong JK, Szigeti-Buck K, Gao Y, Garcia-Caceres C, Yi CX, Salmaso N, Vaccarino FM, Chowen J, Diano S, Dietrich MO, Tschöp MH, Horvath TL. Leptin signaling in astrocytes regulates hypothalamic neuronal circuits and feeding. Nat Neurosci. 2014 Jul;17(7):908–10. doi: 10.1038/nn.3725. Epub 2014 Jun 1. PMID: 24880214; PMCID: PMC4113214.

19. García-Cáceres C, Quarta C, Varela L, Gao Y, Gruber T, Legutko B, Jastroch M, Johansson P, Ninkovic J, Yi CX, Le Thuc O, Szigeti-Buck K, Cai W, Meyer CW, Pfluger PT, Fernandez AM, Luquet S, Woods SC, Torres-Alemán I, Kahn CR, Götz M, Horvath TL, Tschöp MH. Astrocytic Insulin Signaling Couples Brain Glucose Uptake with Nutrient Availability. Cell. 2016 Aug 11;166(4):867–880. doi: 10.1016/j.cell.2016.07.028. PMID: 27518562; PMCID: PMC8961449.

20. Sofroniew MV, Vinters HV. Astrocytes: biology and pathology. Acta Neuropathol. 2010 Jan;119(1):7–35. doi: 10.1007/s00401-009-0619-8. Epub 2009 Dec 10. PMID: 20012068; PMCID: PMC2799634.

21. Puli L, Pomeshchik Y, Olas K, Malm T, Koistinaho J, Tanila H. Effects of human intravenous immunoglobulin on amyloid pathology and neuroinflammation in a mouse model of Alzheimer’s disease. J Neuroinflammation. 2012 May 29;9:105. doi: 10.1186/1742-2094-9-105. PMID: 22642812; PMCID: PMC3416679.

22. Stępień M, Stępień A, Wlazeł RN, Paradowski M, Banach M, Rysz J. Obesity indices and inflammatory markers in obese non-diabetic normo- and hypertensive patients: a comparative pilot study. Lipids Health Dis. 2014 Feb 8;13:29. doi: 10.1186/1476-511X-13-29. PMID: 24507240; PMCID: PMC3921991.

23. Popko K, Gorska E, Stelmaszczyk-Emmel A, Plywaczewski R, Stoklosa A, Gorecka D, Pyrzak B, Demkow U. Proinflammatory cytokines Il-6 and TNF-α and the development of inflammation in obese subjects. Eur J Med Res. 2010 Nov 4;15 Suppl 2(Suppl 2):120–2. doi: 10.1186/2047-783x-15-s2-120. PMID: 21147638; PMCID: PMC4360270.

24. Zhang J, Guo S, Li J, Bao W, Zhang P, Huang Y, Ling P, Wang Y, Zhao Q. Effects of high-fat diet-induced adipokines and cytokines on colorectal cancer development. FEBS Open Bio. 2019 Dec;9(12):2117–2125. doi: 10.1002/2211-5463.12751. Epub 2019 Nov 18. PMID: 31665829; PMCID: PMC6886304.

25. Allen NJ, Eroglu C. Cell Biology of Astrocyte-Synapse Interactions. Neuron. 2017 Nov 1;96(3):697–708. doi: 10.1016/j.neuron.2017.09.056. PMID: 29096081; PMCID: PMC5687890.

26. Matsumura S, Miyakita M, Miyamori H, Kyo S, Shima D, Yokokawa T, Ishikawa F, Sasaki T, Jinno T, Tanaka J, Goto T, Momma K, Ishihara K, Berdeaux R, Inoue K. Stimulation of Gs signaling in MC4R cells by DREADD increases energy expenditure, suppresses food intake, and increases locomotor activity in mice. Am J Physiol Endocrinol Metab. 2022 May 1;322(5):E436–E445. doi: 10.1152/ajpendo.00439.2021. Epub 2022 Mar 28. PMID: 35344393.

27. Griffin JM, Fackelmeier B, Fong DM, Mouravlev A, Young D, O’Carroll SJ. Astrocyte-selective AAV gene therapy through the endogenous GFAP promoter results in robust transduction in the rat spinal cord following injury. Gene Ther. 2019 May;26(5):198–210. doi: 10.1038/s41434-019-0075-6. Epub 2019 Apr 8. PMID: 30962538; PMCID: PMC6760677.

28. Douglass JD, Dorfman MD, Fasnacht R, Shaffer LD, Thaler JP. Astrocyte IKKβ/NF-κB signaling is required for diet-induced obesity and hypothalamic inflammation. Mol Metab. 2017 Jan 28;6(4):366–373. doi: 10.1016/j.molmet.2017.01.010. PMID: 28377875; PMCID: PMC5369266.

29. Zhang X, Zhang G, Zhang H, Karin M, Bai H, Cai D. Hypothalamic IKKbeta/NF-kappaB and ER stress link overnutrition to energy imbalance and obesity. Cell. 2008 Oct 3;135(1):61–73. doi: 10.1016/j.cell.2008.07.043. PMID: 18854155; PMCID: PMC2586330.

30. Xu B, Goulding EH, Zang K, Cepoi D, Cone RD, Jones KR, Tecott LH, Reichardt LF. Brain-derived neurotrophic factor regulates energy balance downstream of melanocortin-4 receptor. Nat Neurosci. 2003 Jul;6(7):736–42. doi: 10.1038/nn1073. PMID: 12796784; PMCID: PMC2710100.

31. Mirza S, Hossain M, Mathews C, Martinez P, Pino P, Gay JL, Rentfro A, McCormick JB, Fisher-Hoch SP. Type 2-diabetes is associated with elevated levels of TNF-alpha, IL-6 and adiponectin and low levels of leptin in a population of Mexican Americans: a cross-sectional study. Cytokine. 2012 Jan;57(1):136–42. doi: 10.1016/j.cyto.2011.09.029. Epub 2011 Oct 28. PMID: 22035595; PMCID: PMC3270578.

32. Altavilla D, Cainazzo MM, Squadrito F, Guarini S, Bertolini A, Bazzani C. Tumor necrosis factor-alpha as a target of melanocortins in haemorrhagic shock, in the anesthetized rat. Br J Pharmacol. 1998 Aug;124(8):1587–90. doi: 10.1038/sj.bjp.0702038. PMID: 9756372; PMCID: PMC1565580.

33. Uysal KT, Wiesbrock SM, Marino MW, Hotamisligil GS. Protection from obesity-induced insulin resistance in mice lacking TNF-alpha function. Nature. 1997 Oct 9;389(6651):610–4. doi: 10.1038/39335. PMID: 9335502.

34. Mina AI, LeClair RA, LeClair KB, Cohen DE, Lantier L, Banks AS. CalR: A Web-Based Analysis Tool for Indirect Calorimetry Experiments. Cell Metab. 2018 Oct 2;28(4):656-666.e1. doi: 10.1016/j.cmet.2018.06.019. Epub 2018 Jul 12. PMID: 30017358; PMCID: PMC6170709.

35. Voss-Andreae A, Murphy JG, Ellacott KL, Stuart RC, Nillni EA, Cone RD, Fan W. Role of the central melanocortin circuitry in adaptive thermogenesis of brown adipose tissue. Endocrinology. 2007 Apr;148(4):1550–60. doi: 10.1210/en.2006-1389. Epub 2006 Dec 28. PMID: 17194736.

36. Stanford KI, Middelbeek RJ, Townsend KL, An D, Nygaard EB, Hitchcox KM, Markan KR, Nakano K, Hirshman MF, Tseng YH, Goodyear LJ. Brown adipose tissue regulates glucose homeostasis and insulin sensitivity. J Clin Invest. 2013 Jan;123(1):215–23. doi: 10.1172/JCI62308. Epub 2012 Dec 10. PMID: 23221344; PMCID: PMC3533266.

37. Alcalá M, Calderon-Dominguez M, Serra D, Herrero L, Viana M. Mechanisms of Impaired Brown Adipose Tissue Recruitment in Obesity. Front Physiol. 2019 Feb 13;10:94. doi: 10.3389/fphys.2019.00094. PMID: 30814954; PMCID: PMC6381290.

