## Supplemental Fig 1 for "Hypothalamic melanocortin-4 receptors on astrocytes mediate inflammation and body weight homeostasis"

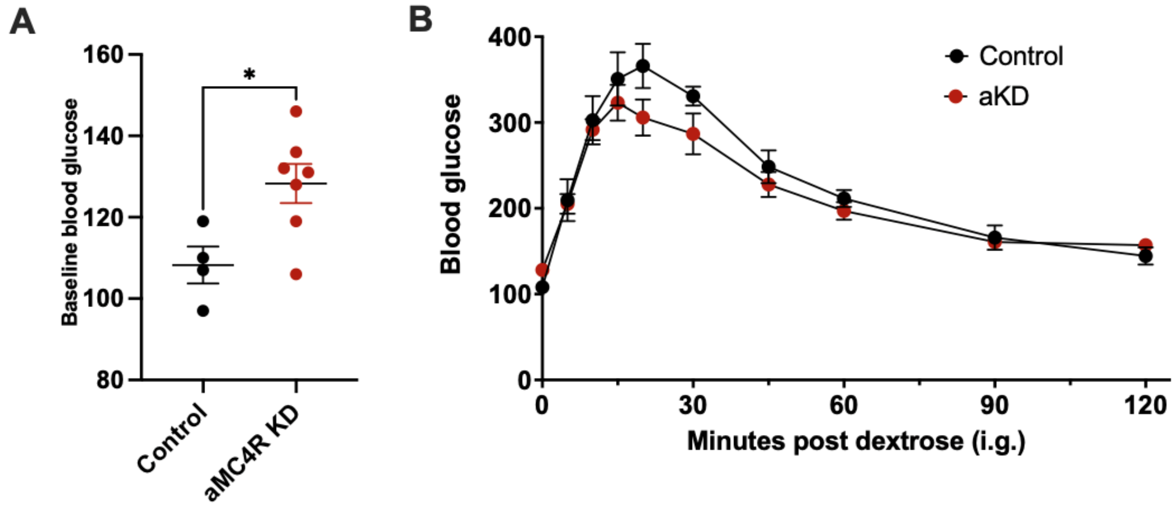


**Supplementary Figure 1. Comparison of blood glucose and glucose tolerance in control and aMC4R KD mice.** Glucose analysis revealed significantly higher fasted blood glucose levels in experimental treated mice *P=0.0228 when compared to controls (Panel A). Oral glucose tolerance test (OGTT) showed no difference between groups with the experimental treated mice exhibiting slightly improved glucose clearance (Panel B).
